# Strontium isotopes reveal diverse life history variations, migration patterns, and habitat use for Broad Whitefish (*Coregonus nasus*) in Arctic, Alaska

**DOI:** 10.1101/2021.11.02.466917

**Authors:** Jason C. Leppi, Daniel J. Rinella, Mark S. Wipfli, Randy J. Brown, Karen J. Spaleta, Matthew S. Whitman

## Abstract

Conservation of Arctic fish species is challenging partly due to our limited ability to track fish through time and space, which constrains our understanding of life history diversity and lifelong habitat use. Broad Whitefish (*Coregonus nasus*) is an important subsistence species for Alaska’s Arctic Indigenous communities, yet little is known about life history diversity, migration patterns, and freshwater habitat use. Using laser ablation Sr isotope otolith microchemistry, we analyzed Colville River Broad Whitefish ^87^Sr/^86^Sr chronologies (n = 61) to reconstruct movements and habitat use across the lives of individual fish. We found evidence of at least six life history types, including three anadromous types, one semi-anadromous type, and two nonanadromous types. Anadromous life history types comprised a large proportion of individuals sampled (collectively, 59%) and most of these (59%) migrated to sea between ages 0–2 and spent varying durations at sea. The semi-anadromous life history type comprised 28% of samples and entered marine habitat as larvae. Nonanadromous life history types comprised the remainder (collectively, 13%). Otolith ^87^Sr/^86^Sr data from juvenile and adult freshwater stages suggest that habitat use changed in association with age, seasons, and life history strategies. This information on Broad Whitefish life histories and habitat use across time and space will help managers and conservation planners better understand the risks of anthropogenic impacts and help conserve this vital subsistence resource.

## Introduction

Global freshwater biodiversity is in decline and many fish species are at risk of extinction [1–4]. Freshwater ecosystems cover less than 1% of the global surface but support about half of the 30,000 described fish species [5]. Consequently, environmental impacts on freshwater ecosystems can have a disproportionate effect on interspecific and intraspecific fish diversity [6–8]. Arctic freshwater ecosystems are no exception and recent assessments confirm that ongoing climate-driven changes [9] threaten Arctic freshwater biodiversity [10].

The accelerated impacts of climate change at high latitudes [11,12] are a major threat to Arctic freshwater ecosystems [13], altering streamflow [14–17], warming [18,19] and drying [20] aquatic habitats; causing eutrophication [21] and browning of lakes [22,23]; and allowing for northward range expansion of eurythermic species [24]. To a lesser extent, long-range pollution [7], habitat loss and degradation, and flow modification from oil and gas development [25] can occur independently or interact with climatic change to alter Arctic freshwater ecosystems [26]. New regional and global emerging threats, such as invasive species [27], diseases, or algal blooms [28] may add additional stress to ecosystems [7], potentially reducing genetic and life history diversity within populations and further accelerating biodiversity loss [29–31].

Understanding lifelong habitat use for Arctic fishes is challenging due to their high mobility and our limited ability to track fish as they move among a suite of habitats (i.e., foraging, overwintering, spawning) that are geographically dispersed, change over time, and are often temporary [32]. As a response to dispersed resources and constantly changing conditions, Arctic fish populations have developed numerous life history strategies to exploit food resources, seek refugia from harsh environmental conditions [33], and maximize reproductive success and survival [34,35].

Foraging strategies include ontogenetic or seasonal movements to marine habitats [36–39] but, due to extreme winter conditions, fish must leave productive marine habitats and find overwintering refugia in freshwater or brackish habitat to avoid lethal low temperatures [40]. An iteroparous reproductive strategy is typical for fishes that use habitats with high environmental variability [41,42] and, when coupled with a longer life, it hedges against unpredictable conditions [43]. The proximity of safe rearing habitats at or downstream of spawning areas also helps to facilitate juvenile survival [44,45] and, for anadromous fishes, a diversity of age at ocean entry helps to buffer the effects of unfavorable conditions [29]. This diversity in life history facilitates long-term population stability, helping to stabilize ecosystems and buffer populations from environmental perturbations that may reduce habitat quality and increase mortality [46–48].

Broad Whitefish (*Coregonus nasus*), a long-lived migratory species, is an important subsistence resource for Alaska’s Indigenous communities. Little is known about the specifics of habitat use within and across life stages in Arctic populations, but previous research in other parts of its range supports the theory of a highly mobile species that utilizes various aquatic habitats [50–52]. Broad Whitefish reach sexual maturity around the age of eight but can live 30 or more years [49]. Habitat use across time and space likely results in a variety of life histories with varying amounts of time spent in freshwater, estuarine, and marine habitats [38,53]. Broad Whitefish larvae are thought to be passively advected downstream to deltas, estuaries, and nearshore areas by spring breakup flows, based on their hatch timing, inability to resist spring streamflow as larvae, and general abundance in coastal and estuarine habitat [54–57]. Referred to as *Aanaakliq* in the Iñupiaq language, Broad Whitefish, are valued due to their relatively large size (up to 4.5 kg) and abundance during migrations [58,59], accounting for about half the total mass of fishes harvested across all Beaufort Sea communities [60]. However, without information on life history diversity and habitat use, land and resource managers lack essential information for the management and conservation of this important subsistence fish.

Otolith microchemistry is an effective tool to understand life history diversity and habitat use of fishes [51,61–66]. Otoliths, paired inner ear stones used for hearing and balance in all teleost fishes, are laid as concentric layers of metabolically inert biogenic minerals, primarily calcium carbonate. Elements are permanently incorporated into their organic matrix, and compositional changes across the layers reflect changes across an individual’s life [67]. Strontium (Sr), a naturally occurring element derived from geologic material, has four stable isotopes (^88^Sr, ^87^Sr, ^86^Sr, ^84^Sr), in which only ^87^Sr is radiogenic. The ratio of ^87^Sr to ^86^Sr (^87^Sr/^86^Sr), which roughly has similar proportions of elemental Sr and can be measured at similar precisions, reflects Sr released into freshwater sources (e.g., rivers, lakes, streams) and is driven by differences in lithology, age, chemical composition [68–70], and weathering rates of surficial geology [71–73].

Fish take up dissolved Sr through the gills, and the Sr isotopes are incorporated into the otolith matrix [67,74], forming a continuous record of Sr isotope values that directly reflects habitats inhabited across an individuals’ life[75]. Compared to freshwater habitats, ^87^Sr/^86^Sr in marine habitats are generally lower, homogenous, and constant due to the long residence time and mixing of oceans [71]. The concentrations of total Sr and ^88^Sr, the most abundant of the four stable isotopes and a reliable proxy for total Sr, increase with water salinity and are consistently higher in marine versus freshwater habitats. For diadromous fishes, these relative differences in Sr isotopes can be used to indicate the timing and duration of freshwater, estuarine, and marine habitat use [38,76].

Broad Whitefish populations use the Colville River watershed for foraging [77], rearing [77], and spawning [78]. With headwaters in the rugged Brooks Range, the Colville River is one of the few rivers in the region that contains abundant gravel substrate and deep channels, which are both likely essential for egg survival [49]. Due in part to its watershed size, the Colville River also has the largest delta on the Alaskan Beaufort Sea coast, which provides abundant rearing habitat for larval and juvenile fishes [79]. Broad Whitefish can live for 30+ years, and they return to the Colville River ecosystem regularly to reproduce [59], likely migrating from a variety of productive foraging areas in coastal lagoons and rivers across the Beaufort Coastal Plain. Thus, by sampling the Colville River’s spawning run, we may be learning about life-history patterns of Broad Whitefish at the regional scale.

Conservation of freshwater fish diversity requires an understanding of lifelong habitat use. For many Arctic fishes, this information is lacking because movement between key habitats changes with age and life history strategy and is difficult to monitor. To fill important knowledge gaps about Broad Whitefish habitat use, we used Sr isotope (^87^Sr/^86^Sr, ^88^Sr) otolith chronologies across individuals’ lives to quantify life history attributes and reconstruct migration patterns of migrating fish captured within the Colville River, Alaska. Our specific objectives were to (i) document the range of life history types and explore how individuals are distributed among several predefined life history types, (ii) determine the proportion of Broad Whitefish that are anadromous and investigate the timing of marine habitat use, and (iii) determine if freshwater natal and freshwater juvenile rearing region ^87^Sr/^86^Sr remain constant across life history groups. Understanding the diversity of lifelong habitat use will provide managers and conservation planners essential information to begin to understand Broad Whitefish exposure to changing habitats.

## Materials and Methods

### Study area

The Central Beaufort Sea region study area (Fig 1) contains a diversity of fish habitats. Situated between the Ikpikpuk and Canning rivers in Arctic Alaska, the coastline is a spectrum of bays and inlets, tapped basins (basins that are breached by the sea due to erosion), lagoons behind barrier islands, and exposed bluffs [80]. River deltas are frequent along the coast, the size varying based on physiography and watershed size [80]. Thermokarst and riverine lakes that vary in size, depth, and connectivity [81,82] cover 30% of the region’s surface area [83]. Stream habitats vary by watershed and geomorphic setting [84,85], resulting in colluvial channels in foothill and mountainous headwaters, beaded headwater streams in low-gradient coastal plains, and meandering alluvial streams and rivers lower in watersheds [82]. Geologic lithology consists of materials from Precambrian and Paleozoic fragments and continental margin materials dating to the Cretaceous period [68]. Modeled ^87^Sr/^86^Sr are estimated to be highest in the Brooks Range (ca. 0.7200), which contains sedimentary noncarbonate lithology with minor areas of intrusive mafic and ultramafic lithologies, and lowest near the coast (ca. 0.7070) within marine and eolian deposits [69]. Modeled stream ^87^Sr/^86^Sr tend to be higher in watersheds that drain high relief areas of the Brooks Range but also remain relatively high in the mainstem Colville River until the tidally influenced areas of the lower river [68,69]. Modeled stream ^87^Sr/^86^Sr use bedrock and chemical weathering models within a streamflow accumulation model to estimate values at 1 km grid cell, which does not account for modification of isotope values during transport [86].

**Fig 1.**
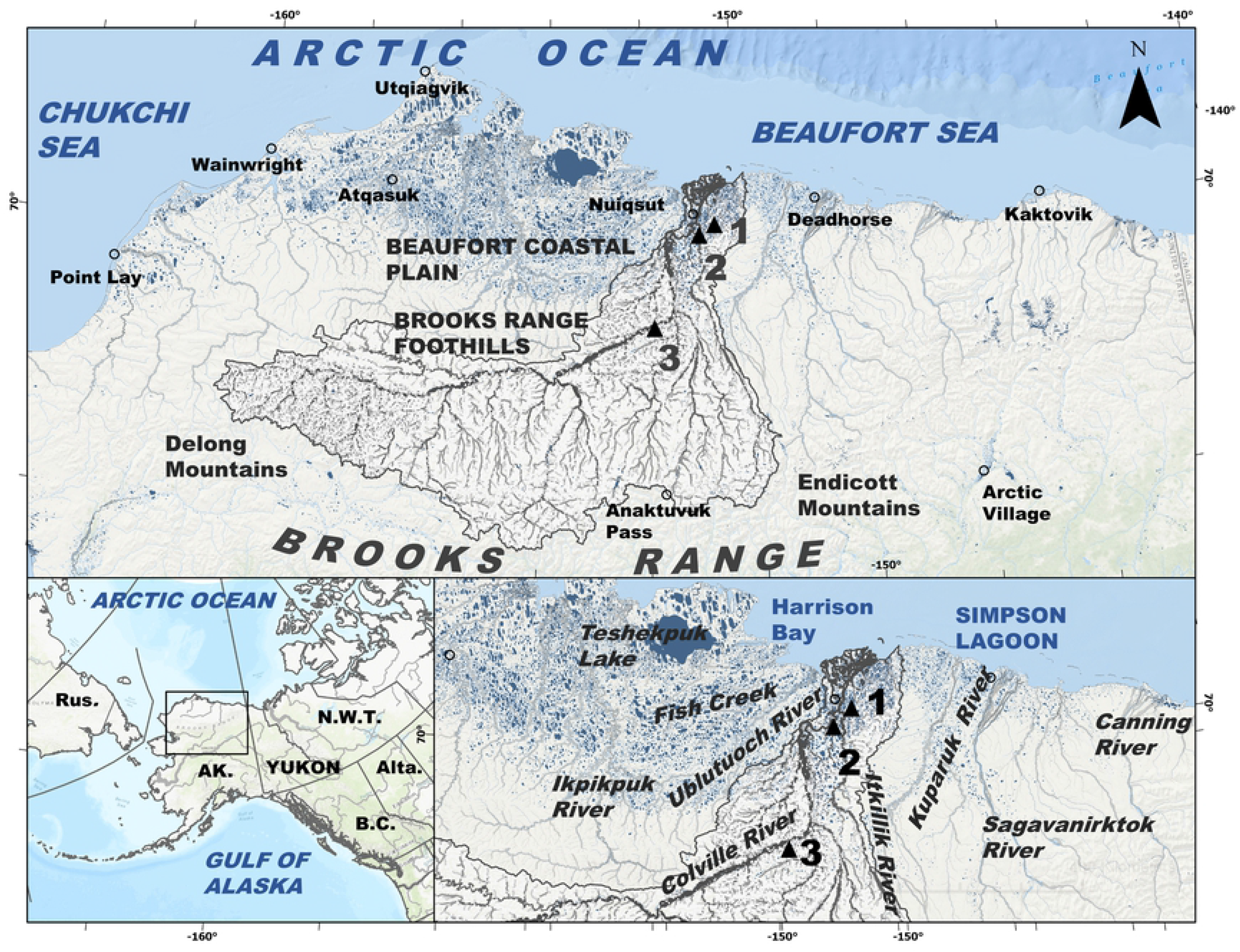
Study area. The Central Beaufort Sea region in Arctic Alaska, situated between the Ikpikpuk and the Canning River, AK, contains a diversity of aquatic habitats. The large Colville River, AK, USA (ca. watershed area 60,000 km^2^), located in the middle of the Central Beaufort Sea coast, AK, contains minor tributaries that drain from the Brooks Range, AK (thin grey lines) and main tributaries (thick dark grey lines) that flow toward a large delta on the edge of the Beaufort Sea, near the community of Nuiqsut, AK. We collected fish at three sites (black triangles) within the Colville River (site 1= Itkillik, site 2 = Puviksuk, site 3 = Umiat).

The Central Beaufort Sea region, within the Arctic tundra biome, is characterized by permafrost, extreme climate, low-growing plants, and large seasonal variations in day length. The region’s stark seasonality can be divided into two main seasons, cold and warm, but the former controls many of the physical and biological processes. Cold season air temperatures are consistently well below freezing, creating a landscape dominated by snow and ice for about eight months [87]. The warm season is brief but, with 24 hours of daylight and moderate air temperatures [87], the area becomes productive foraging and rearing habitat for many resident and migratory fishes and other animals. Annual precipitation is generally low, with more falling in the foothills than along the coast (30 and 20 cm, respectively) and about half falling as snow [87].

### Fish sampling and otolith collection

We collected sagittal otoliths from adult Broad Whitefish migrating up the Colville River (Fig 2A) during 2015. The Colville, the largest river in Arctic Alaska, flows about 560 km northward from its headwaters in the partially glaciated Brooks Range to a large delta on the edge of the Central Beaufort Sea coast, near the Native Village of Nuiqsut (Fig 1). We set gill nets ca. 30 m in length composed of braided nylon and monofilament with 10-cm and 12-cm stretched mesh to target adult fish large enough to spawn (> 35 cm; [49,88]). We positioned nets at gravel point bars, along eddy lines, and perpendicular to flow in low-gradient reaches at three separate sites (Sites 1–3; Fig 1). We sampled at Puviksuk on July 23–27, at Umiat on August 21–26, and at Itkillik on October 10–11. We euthanized captured Broad Whitefish and recorded fork length (n = 97, all of which were adults ≥ 42 cm; [49]), total weight, gonad weight, and sex (44 males, 47 females, 7 undetermined; S1 Table). We collected sagittal otoliths from all individuals using the Guillotine method [89], rinsed in water, and stored in paper envelopes (Table 1). The planned sample size of 50 individuals per site was smaller than anticipated at Umiat (n = 23) and Itkillik (n = 17), as opposed to Puviksuk (n = 57), due to unexpectedly high streamflow at the former and an early freeze-up at the latter that inhibited our ability to capture fish.

**Table 1.**
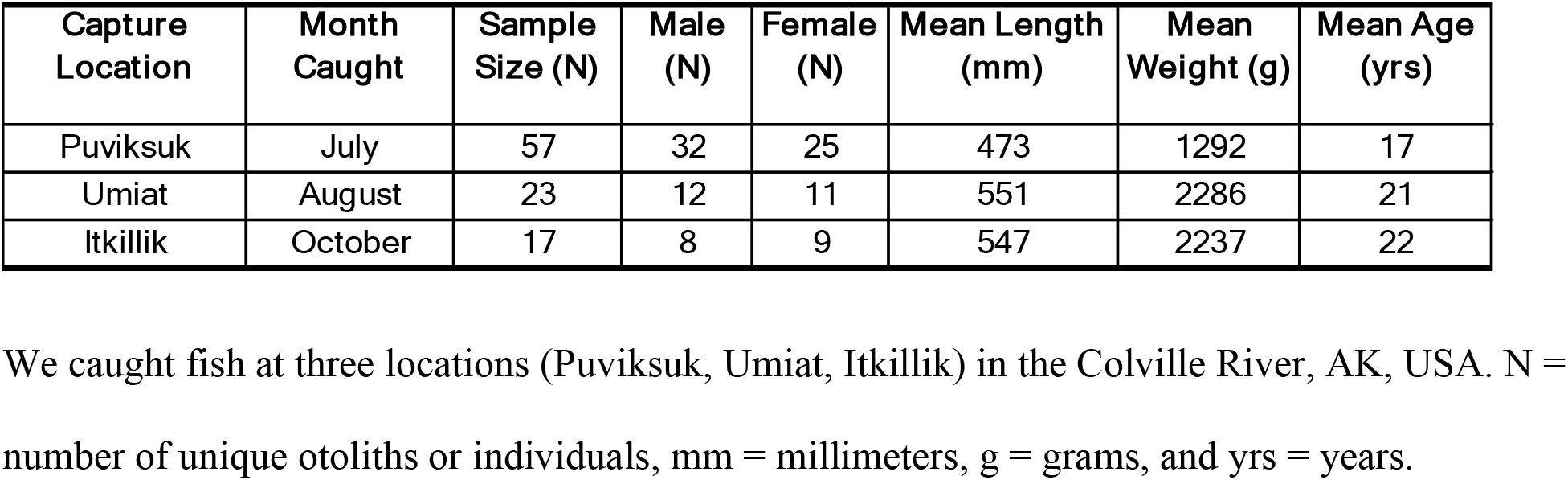
Summary of otoliths collected from Broad Whitefish (*Coregonus nasus*) caught in the Colville River, AK, USA.

| Capture Location | Month Caught | Sample Size (N) | Male (N) | Female (N) | Mean Length (mm) | Mean Weight (g) | Mean Age (yrs) |
| --- | --- | --- | --- | --- | --- | --- | --- |
| Puviksuk | July | 57 | 32 | 25 | 473 | 1292 | 17 |
| Umiat | August | 23 | 12 | 11 | 551 | 2286 | 21 |
| Itkillik | October | 17 | 8 | 9 | 547 | 2237 | 22 |

**Fig 2.**
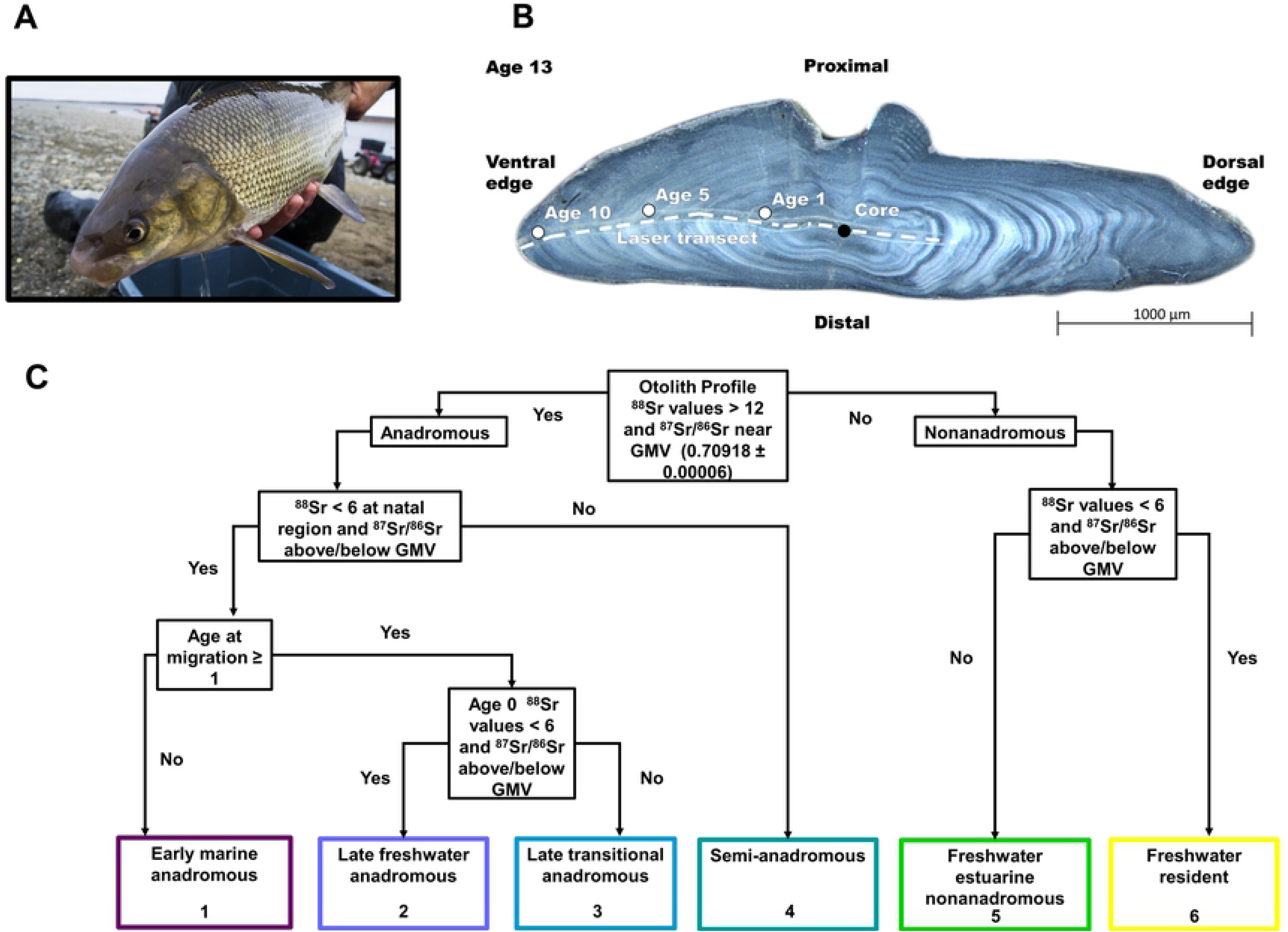
Otolith collection, laser analysis, and life history type classification. (A) Broad Whitefish (*Coregonus nasus*) from the Colville River, AK, USA (Photo credit: Jason C. Leppi) (B) Broad Whitefish otolith showing the core, annuli, and laser ablation path (Photo credit: Dan Bogan). (C) Flow diagram showing the life history supervised classification approach used to group Broad Whitefish.

We caught fish at three locations (Puviksuk, Umiat, Itkillik) in the Colville River, AK, USA. N = number of unique otoliths or individuals, mm = millimeters, g = grams, and yrs = years.

### Sr isotope analysis of otoliths

#### Otolith preparation

We mounted sagittal otoliths on individual microscope slides using Crystalbond^™^ 509, smoothed using a 1200-grit grinding wheel, and then thin sectioned in the transverse plane. We then polished thin-sectioned otoliths with 3-μm alumina slurry until the core was visible [90], photographed, and visually aged using standard methods [91] and an Olympus^™^ microscope with MicroPublisher^™^ imaging attachment. In preparation for isotope analysis, we remounted otoliths on petrographic slides (2.7 x 4.6 cm) and repolished using 3-μm alumina slurry. A total of five petrographic slides held all 97 otoliths and each otolith was tracked and labeled using a numerical ID.

#### Laser ablation

We measured Sr isotope concentrations (^88^Sr, ^87^Sr, ^86^Sr, ^84^Sr) across a subset of the otoliths (69 of 98 individuals sampled) due to instrument constraints (i.e., time, funding, instrument availability). We selected samples to prioritize for laser ablation with the goal of analyzing otoliths across a gradient of δ^13^Cˈ, δD, and δ^18^O tissue values, which was expected to represent time spent in different habitat types (e.g., freshwater, estuarine, marine) over the three months prior to capture (S3 Table; [78]). Subsampled otoliths were roughly proportional to the number of samples collected at each field site [78].

We used an Analyte G2 Excimer 193-nm Laser Ablation System (LA; Teledyne Photon Machines, Bozeman, USA) with a Helex cell coupled to a Neptune Plus^™^ multi-collector inductively coupled plasma mass spectrometer (MC-ICP-MS; Thermo Scientific^™^, Bremen, Germany) for strontium isotope analyses at the University of Alaska Fairbanks’ Alaska Stable Isotope Facility. We selected an ablation path perpendicular to annuli across each otolith and, to ensure that the freshwater natal region was identified, our ablation path encompassed the entire first year of growth (Fig 2B) [63]. We performed an initial ablation cleaning using 80 μm circle spot size and 15 μm/s scan speed over the surface of otoliths prior to analysis. For isotope analysis, the laser was set to 35 μm beam diameter, a pulse rate of 10 Hz, a speed of 5 μm/s, and a laser fluence of 6.64 J/cm^2^. We mixed the laser stream with the output of an Aridus II^™^ membrane desolvation system via a mixing chamber immediately prior to the plasma inlet, which allowed for optimization of the signal of the MC-ICP-MS and facilitated external intra-elemental correction of the instrument isotope fractionation (IIF) [92]. We used a National Institute of Standards and Technology (NIST^®^) solution of ~13 ng g^−1^ SRM 987 (SrCO_3_ isotopic standard) and ~20 ng g^−1^ Zr standard to optimize the instrument and also aspirated via the Aridus II^™^ every 30 minutes for IIF correction via bracketing from standard to sample acquisition. While we ablated samples, the Aridus II^™^ aspirated a 2% solution of HNO_3_ that was double distilled via sub-boiling distillation (Savillex DST-1000) from trace metal grade concentrated HNO_3_. We then used American Society for Testing and Materials (ASTM) Type I water from a Milli-Q^®^ IQ 7000 system, which had been further purified via a double sub-boiling distillation, for dilution. Instrumental parameters used during laser ablation isotopic analysis are listed in the S2 Table of the Appendix.

#### Data reduction

We processed ablation data as outlined in Irrgeher (2016) [95]. We blank-corrected data by subtracting the mean gas blank values obtained during the 30-second laser-warmup period prior to each ablation for all measured isotopes. We calculated calcium (Ca) dimer (CaCa) and Ca argide (CaAr) corrections based on the blank corrected signals at mass to charge number (m/z = 82 and 83) using the natural abundances of Ca and argon (Ar) and applied to the blank corrected signals at m/z 84, 85, 86, 87, and 88. We calculated rubidium (Rb) interference correction on m/z 87 using Russell’s law via mass 85 and applied via peak stripping in which the fractionation of Rb was assumed to be equal to strontium (Sr) due to the low levels of Rb naturally occurring in sample materials like otoliths. We corrected the ^87^Sr/ ^86^Sr ratio via standard/sample bracketing of the NIST SRM^®^987 solution. Instrumental isotopic fractionation (IIF) was corrected externally, also using the NIST SRM^®^987 solution.

#### Otolith ^87^Sr/ ^86^Sr data post-processing

We individually inspected strontium data from otolith laser ablation for data quality, trimmed, and adjusted accordingly. We removed otoliths from the dataset if Sr data contained unreliable values due to cracks (n = 8). We used a Leica^®^ microscope with a micrometer to measure the length of the ablation path from age-1 dorsal side to the otolith edge on the ventral side (Fig 2B). We visually identified otolith core and annuli following standard methods [91] and measured the length of each annuli from the beginning of the ablation transect using a Leica^®^ microscope with a micrometer. We then cropped cropped data in R statistical program (http://cran.r-project.org/), from the estimated otolith core to the edge, to facilitate comparison between otoliths. Finally, to facilitate classification we inspected strontium data for each otolith and made minor adjustments to the ratio data (^87^Sr/^86^Sr) to align them with the global marine value (GMV = 0.70918 ± 0.00006) when ^88^Sr concentrations were above 12.26 V and ensure that non-elevated ^88^Sr concentrations (< 12.26 V) fell outside of this range.

#### Statistical analysis

We used Generalized Additive Models (GAMs) to analyze the ^87^Sr/^86^Sr variation from each otolith core to edge. A GAM is an extension of a generalized linear model in which the linear predictor is provided by a user-assigned sum of smooth functions of the covariates, plus a conventional parametric component of the linear predictor [96]. We fit GAMs to each otolith ^87^Sr/^86^Sr profile using the MGCV package in R, which is similar to previous methods used to analyze otolith ^87^Sr/^86^Sr of Slimy Sculpin (*Cottus cognatus*) [97] and Chinook salmon (*Oncorhynchus tshawytscha*) [62]. MGCV implementation of GAMs uses penalized iteratively re-weighted least squares (P-IRLS) to maximize goodness-of-fit, solving the smoothing parameter estimation problem and the Generalized Cross-Validation Criterion to minimize overfitting of the data [96,98]. We determined GAM model parameters through iterative model fit comparisons and for final GAMs we set the lambda (λ) smoothing parameter to 0.6 and knots parameter (k) to 100. Finally, we calculated Bayesian 95% confidence intervals along each ^87^Sr/^86^Sr profile.

### Life history attributes and classification

We visually compared the ^88^Sr concentrations and ^87^Sr/^86^Sr across each otolith core-to-edge chronology and, based on inferred life history patterns, developed a supervised classification approach that grouped individuals into life history types. Initial life history types included anadromous, semi-anadromous, and nonanadromous, but during initial visual inspection of otolith profiles it was apparent that subgroups existed within these groups. We estimated otolith Sr concentrations (Sr mg/kg) by dividing the concentration of Sr in the FEBs-1 standard (i.e., 2055 mg/kg) by the average ^88^Sr FEBs-1 standard value during ablation and multiplying by ^88^Sr otolith values at individual points across the ablation path. We considered a otolith ^88^Sr signal below 6.13 V (corresponding to ca. 850 mg/kg) to be time spent in freshwater habitat, a signal greater than 6.13 V and less than 12.26 V (corresponding to ca.1700 mg/kg) to be time spent in estuarine water and a signal greater than 12.26 V (corresponding to ca.1700 mg/kg) to be marine habitat use [38,76].

We inferred occupied habitats and behaviors based on otolith ^88^Sr concentrations and ^87^Sr/^86^Sr at different ages, determined by reading annuli and identifying important otolith regions. We defined natal regions as areas distal to the core and without detectable maternal strontium that had ^88^Sr concentrations representative of freshwater and relatively constant ^87^Sr/^86^Sr, which were assumed to precede the onset of exogenous feeding and downstream migration to rearing habitat [99]. We defined freshwater juvenile rearing regions as all values distal to the natal region, but before migration to marine habitats and prior to age-1(i.e., freshwater age-0 juvenile rearing period). We considered freshwater regions to be isotopically distinct from each other, inferring movement to a different habitat, if the difference between mean ^87^Sr/^86^Sr was > 0.00005 [97]. This value was chosen because it was sufficiently greater than the mean ± 2 S.E. of all juvenile and natal region values and would be large enough to detect differences [62,97]. For individuals that did not migrate to marine habitat at age-0, this region encompassed all values during the growth period (identified as the opaque region with wide growth bands) before the winter period [91]. We determined age at marine entry by comparing ^88^Sr concentrations and ^87^Sr/^86^Sr in relation to annuli and we assigned all anadromous individuals with marine entry values (e.g., age-0, age-1) based on when the first maximum ^88^Sr concentrations were above 12.26 V and ^87^Sr/^86^Sr were near the GMV.

We grouped individuals into six life history types using a supervised classification approach. We considered individuals with otolith ^88^Sr concentrations above 12.26 V and ^87^Sr/^86^Sr near the GMV anadromous, and we considered all other individuals as nonanadromous [38,51,63]. We classified anadromous individuals into four subgroups based on the type of rearing habitat inhabited at age-0 and duration in fresh water prior to marine entry (Fig 2C). We classified individuals as follows: i) early marine anadromous (Type 1) if age at migration from fresh water was less than one; ii) as late freshwater anadromous (Type 2) if age-0 ^87^Sr/^86^Sr represented time spent in fresh water and age at migration was greater than one; and iii) late transitional anadromous (Type 3) if age zero ^87^Sr/^86^Sr represented time spent in both fresh and estuarine water and age at migration was greater than one (Fig 2C). We classified individuals as semi-anadromous if ^87^Sr/^86^Sr at the natal region was near GMV (Type 4; Fig 2C). Semi-anadromous individuals had no detectable age-0 freshwater otolith signature, likely spending limited time in fresh water as larvae and frequently moving between freshwater, estuarine, and marine habitats (Fig 2C). We classified nonanadromous individuals into two subgroups based on the habitat used throughout their life. We classified individuals as freshwater estuarine nonanadromous (Type 5) if they spent time in fresh and brackish water but ^88^Sr concentrations and ^87^Sr/^86^Sr never indicated they entered marine habitats. The remaining nonanadromous individuals showed no evidence of using brackish habitat and therefore we classified these individuals as freshwater residents (Type 6) (Fig 2).

## Results

### Life history types

Anadromous life history (Types 1–3) comprised a significant proportion of individuals sampled (59%; Fig 3A). Early marine anadromous (Type 1) comprised 21%, late freshwater anadromous (Type 2) comprised 23%, and late transitional anadromous (Type 3) comprised 15% (Fig 3A.). All anadromous individuals spent a portion of time (months to years) in fresh water (Fig 4A–C) and had similar natal (Fig 3B) and juvenile rearing ^87^Sr/^86^Sr (Fig 3C). Following initial migration to marine habitats, our results suggested that numerous anadromous individuals shifted from annual marine habitat use (Fig 5A) to constant freshwater residency (Fig 5B). Semi-anadromous (Type 4) individuals were the most common life history type and comprised 28% of the population sampled (Fig 3A). The majority of these individuals had lower natal (Fig 3B) and juvenile freshwater rearing (Fig 3C) ^87^Sr/^86^Sr reflective of estuarine habitats and ^87^Sr/^86^Sr near the GMV that remained somewhat constant across time (Fig 4D). Nonanadromous individuals (Type 5–6) were comparatively rare (13%; Fig 3A), with freshwater estuarine nonanadromous (Type 5) comprising 3%, and freshwater resident (Type 6) comprising 10% (Fig 3A). These nonanadromous individuals had elevated freshwater natal region ^87^Sr/^86^Sr compared to the other life history groups (Fig 3B) but had similar freshwater juvenile rearing ^87^Sr/^86^Sr (Fig 3C). Freshwater estuarine life history types mainly used habitat with ^87^Sr/^86^Sr within the range of fresh waters, but on occasions these individuals used estuarine habitat (Fig 5C). Within the freshwater resident life history type, some individuals appear to use freshwater habitats across a spectrum of ^87^Sr/^86^Sr (Fig 4F), while others remain in habitat with relatively uniform values (Fig 5D).

**Fig 3.**
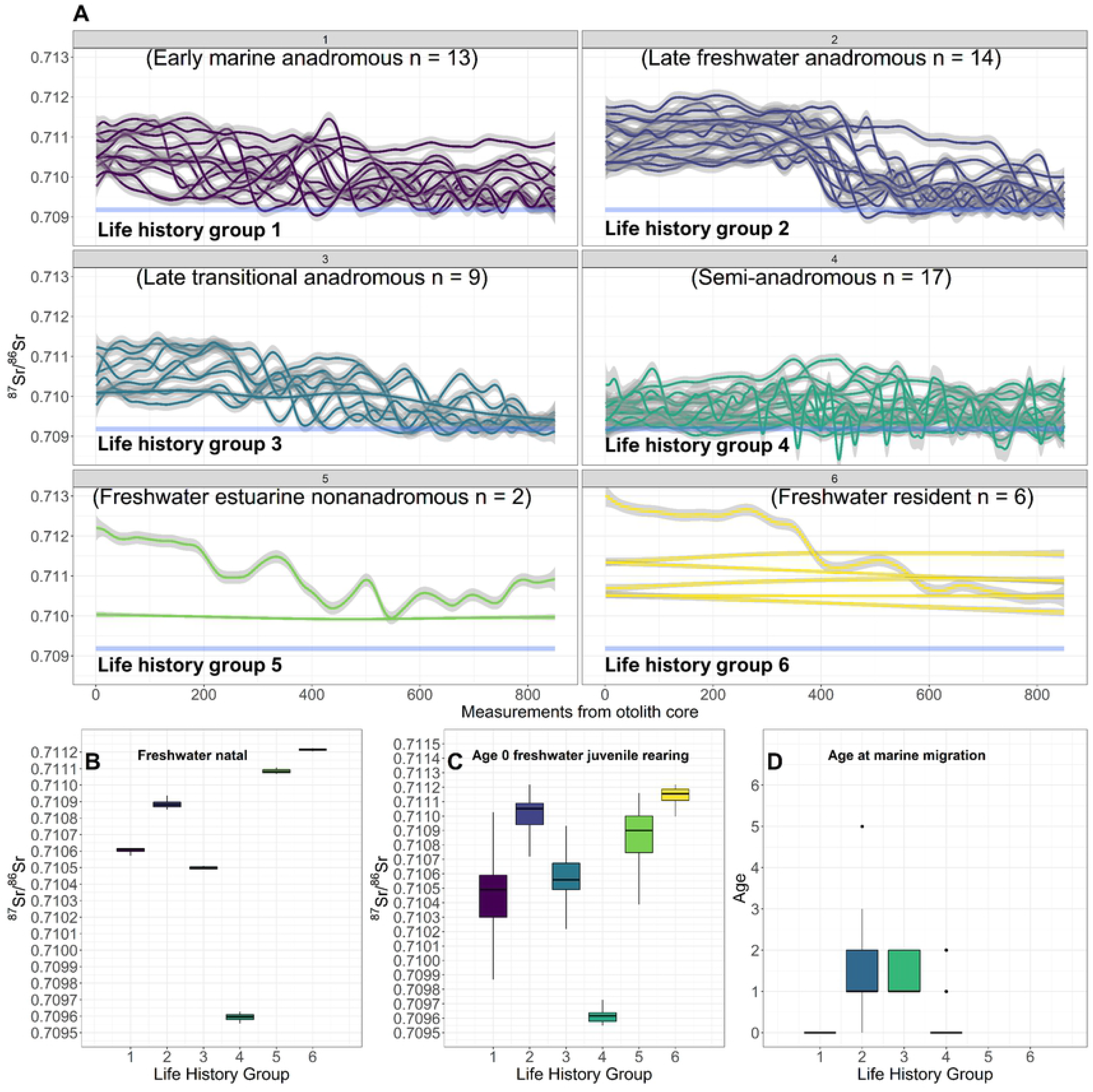
Otolith life history classification. Supervised classification of 61 Broad Whitefish (*Coregonus nasus*) from the Colville River, AK, USA into six life history types based on strategy (anadromous, semi-anadromous, nonanadromous), rearing habitat used (marine, brackish, freshwater), and age at marine migration. Anadromous individuals (Types 1–3), semi-anadromous (Type 4), and non-anadromous (Types 5–6) are individually colored by life history type. Fitted Generalized Additive Model values of the first 850 raw ^87^Sr/^86^Sr (solid colored lines) and 95% confidence interval (grey shading) for each individual and life history type are shown in panel (A). The horizontal light blue line represents the global mean oceanic value (GMV = 0.70918 ± 0.00006). Panel (B) and (C) show boxplots of the freshwater natal region and the freshwater juvenile rearing region. Panel (D) shows boxplots for the age of first marine migration for each life history group. Each boxplot shows the median value (^87^Sr/^86^Sr value or age) for each life history type (horizontal black line), IQR (box outline), the maximum value within 1.5 times the IQR (vertical black line), and outside values greater than 1.5 times the IQR (black dots).

**Fig 4.**
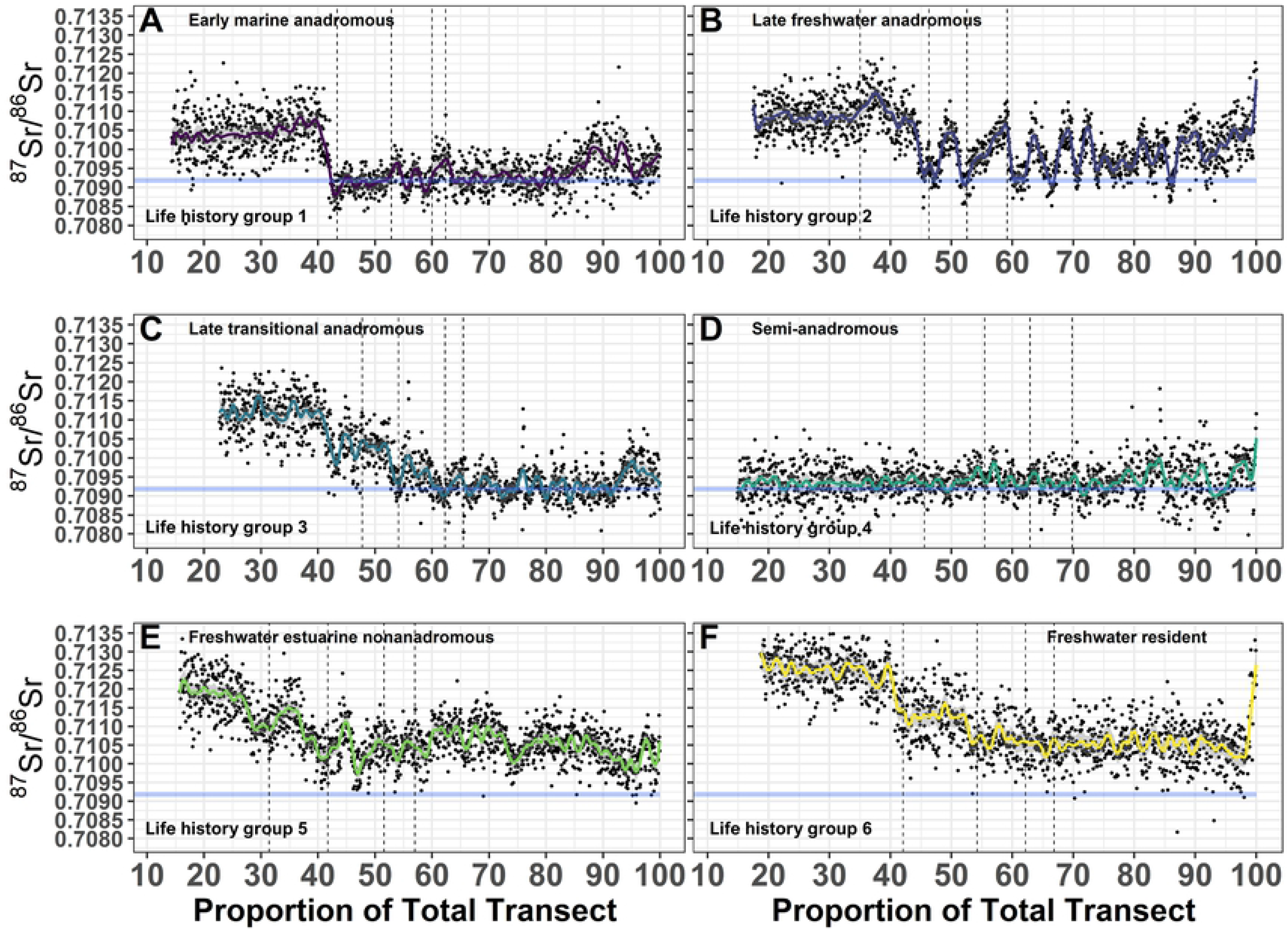
Broad Whitefish life history examples. Selected otolith ^87^Sr/^86^Sr profiles for six Broad Whitefish (*Coregonus nasus*) from the Colville River, AK, USA, representative of each life history type. Fitted Generalize Additive Model values (solid-colored lines), 95% confidence interval (grey shading), and raw ^87^Sr/^86^Sr for each life history type (black dots) from the otolith core to edge are shown. Life history types shown; early marine anadromous (A), late freshwater anadromous (B), late transitional anadromous (C), semi-anadromous (D), freshwater estuarine nonanadromous (E), freshwater resident (F). Thin dashed vertical black lines represent the end of each winter annular growth (annuli) for the first four years. The horizontal light blue line represents the global mean oceanic value (GMV = 0.70918 ± 0.00006).

**Fig 5.**
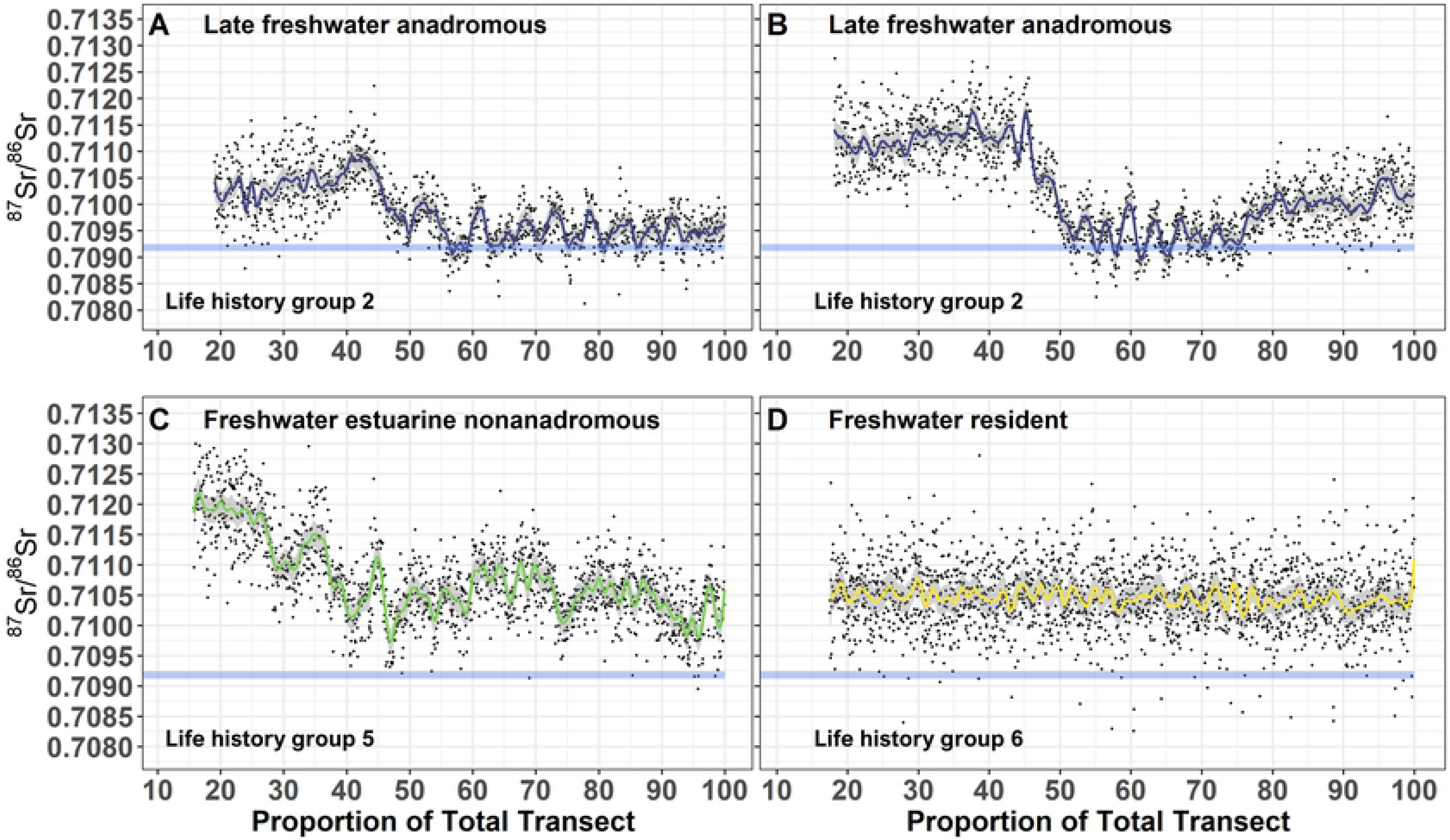
Variation within anadromous and nonanadromous life history types. Variation within anadromous and nonanadromous life history types. Selected otolith ^87^Sr/^86^Sr profiles of four Broad Whitefish (*Coregonus nasus*) from the Colville River, AK, USA, displaying unique variation in anadromous and nonanadromous life history types. Fitted Generalized Additive Model values (solid-colored lines), 95% confidence interval (grey shading), and raw ^87^Sr/^86^Sr for each life history type (black dots) from the otolith core to edge are shown. Freshwater late anadromous life history type displaying continued marine habitat use (A), freshwater late anadromous life history type displaying a switch to freshwater habitat later in life (B), freshwater estuarine nonanadromous life history type demonstrating delayed estuarine habitat use (C), and freshwater resident life history type demonstrating relatively stable ^87^Sr/^86^Sr across time (D). The horizontal light blue line represents the global mean oceanic value (GMV = 0.70918 ± 0.00006).

### Anadromous migration patterns

Among the 59% classified as one of the three anadromous types, most individuals (67%) initially migrated to sea between ages 0 and 2 (Table 2). All anadromous individuals spent months to years in fresh water, but comparisons of GAM values indicate that distinct patterns exist for anadromous types (Fig 3A). Age of marine migration varied based on life history (Table 2), but we found that 39% of individuals migrated to marine habitats at age-0, 25% migrated at age one, 3% migrated at age two, 6% at age three, no fish migrated at age 4, and 3% migrated at age 5 (Fig 3D). Results also show large variation in the duration of marine habitat use (S1 Figure), with some individuals using marine habitats for only a few years, while others consistently use marine habitat until captured (Fig 5A; S1 Figure). Interestingly, our results also show that some adults later in life stop migrating to marine habitats and switch back to using freshwater habitats (Fig 5B; S1 Figure).

**Table 2.** Age at migration for anadromous life history types of Broad Whitefish (*Coregonus nasus*) caught in the Colville River, AK, USA.

| Anadromous Life History Types | Age at Marine Migration |  |  |  |  |  |
| --- | --- | --- | --- | --- | --- | --- |
|  | Age 0 | Age 1 | Age 2 | Age 3 | Age 4 | Age 5 |
| Early marine (Type 1, n = 13) | 100% | 0% | 0% | 0% | 0% | 0% |
| Late Freshwater (Type 2, n = 14) | 0% | 71% | 7% | 14% | 0% | 7% |
| Late Transistional (Type 3, n = 9) | 0% | 56% | 44% | 0% | 0% | 0% |

### Freshwater natal and juvenile rearing region ^87^Sr/^86^Sr

A comparison of freshwater natal and juvenile rearing region ^87^Sr/^86^Sr across life history groups revealed several patterns. Anadromous life history types (early marine, late freshwater, late transitional) had similar values with mean natal region ^87^Sr/^86^Sr ranging from 0.71052 to 0.71083 (Table 3). Individuals with the semi-anadromous life history type had natal region ^87^Sr/^86^Sr centered at 0.70961 and were the lowest values observed (Table 3). Nonanadromous individuals (freshwater estuarine, freshwater resident) had mean natal region ^87^Sr/^86^Sr ranging from 0.71117 to 0.71122 and were the highest values observed (Table 3).

**Table 3.** Summary of GAM predicted ^87^Sr/^86^Sr for freshwater natal and freshwater juvenile rearing regions across life history types of Broad Whitefish captured in the Colville River, AK, USA.

| | $^{87}\text{Sr}/^{86}\text{Sr}$ | | | | | |
| --- | --- | --- | --- | --- | --- | --- |
|  | Mean | Median | Min | Max | SE | N |
| **Freshwater natal region** |  |  |  |  |  |  |
| Early marine anadromous | 0.71058 | 0.71063 | 0.71025 | 0.71083 | 0.000023 | 12 |
| Late freshwater anadromous | 0.71083 | 0.71088 | 0.71036 | 0.71097 | 0.000022 | 14 |
| Late transitional anadromous | 0.71052 | 0.71051 | 0.71020 | 0.71084 | 0.000018 | 9 |
| Semi-anadromous | 0.70961 | 0.70960 | 0.70958 | 0.70986 | 0.000011 | 6 |
| Freshwater estuarine nonanadromous | 0.71106 | 0.71108 | 0.71055 | 0.71114 | 0.000026 | 2 |
| Freshwater resident | 0.71117 | 0.71122 | 0.71062 | 0.71122 | 0.000020 | 6 |
| **Freshwater juvenile rearing region** |  |  |  |  |  |  |
| Early marine anadromous | 0.71048 | 0.71050 | 0.70997 | 0.71091 | 0.000016 | 12 |
| Late freshwater anadromous | 0.71092 | 0.71105 | 0.70981 | 0.71114 | 0.000020 | 14 |
| Late transitional anadromous | 0.71056 | 0.71057 | 0.70992 | 0.71104 | 0.000009 | 8 |
| Semi-anadromous | 0.70966 | 0.70962 | 0.70957 | 0.71008 | 0.000013 | 7 |
| Freshwater estuarine nonanadromous | 0.71087 | 0.71090 | 0.71002 | 0.71113 | 0.000012 | 2 |
| Freshwater resident | 0.71108 | 0.71115 | 0.70982 | 0.71120 | 0.000013 | 6 |

Mean = mean $^{87}\text{Sr}/^{86}\text{Sr}$ , Median = median $^{87}\text{Sr}/^{86}\text{Sr}$ , Min = minimum $^{87}\text{Sr}/^{86}\text{Sr}$ , Max = maximum $^{87}\text{Sr}/^{86}\text{Sr}$ , SE = standard error of $^{87}\text{Sr}/^{86}\text{Sr}$ , and N = number of unique otoliths within the classified life history type.

Freshwater juvenile rearing ^87^Sr/^86^Sr remained relatively constant for some life history types, while others varied. Freshwater juvenile rearing ^87^Sr/^86^Sr for anadromous life history types remained similar (range: 0.71048–0.71092), but early marine anadromous life history type values decreased slightly (−0.00010), while late freshwater anadromous and late transitional anadromous ^87^Sr/^86^Sr increased (+0.00009, +0.00004; Table3). Semi-anadromous life history type ^87^Sr/^86^Sr remained at the lowest values, but freshwater juvenile rearing ^87^Sr/^86^Sr increased from natal values (+0.00005; Table 3). Freshwater estuarine nonanadromous life history type ^87^Sr/^86^Sr decreased by −0.00018 to a mean value of 0.71087, and freshwater resident life history type ^87^Sr/^86^Sr decreased slightly (−0.00009) to 0.71108 (Table 3).

## Discussion

Otolith microchemistry revealed a diversity of life histories in Broad Whitefish in the Colville River. Six different ^87^Sr/^86^Sr life history types were evident in the individuals examined, which included three anadromous, one semi-anadromous, and two nonanadromous life history types. Within each life history group, there also appears to be variation within freshwater habitats occupied during juvenile and adult stages and the time and duration spent in marine habitats. The majority of Broad Whitefish had anadromous life history types, but migration timing varied greatly within life history types. These results support the pattern of increased life history diversity among species that live in environments subjected to frequent perturbations and extreme seasonal changes (e.g., shifting river channels, river ice growth up to 2 m thick), which can increase mortality risks [100].

### Broad Whitefish life history diversity

The observed diversity of life histories and habitats utilized by Broad Whitefish in this study is generally consistent with other studies conducted on coregonins in Arctic and Boreal regions [38,51,63,101]. Research in the Mackenzie River watershed, in Yukon, Canada, showed multiple Broad Whitefish life histories [102], including anadromous, semi-anadromous, and nonanadromous (riverine and lacustrine) forms [38,53,103], with large variability in migration patterns and utilization of the watershed [104]. High variation in the timing of marine entry (years) by anadromous individuals was observed in the Mackenzie River, along with a high degree of variability in freshwater, estuarine, and marine habitat use [38]. In the Yukon River, Alaska, otolith Sr concentrations suggest a slow, gradual downstream movement by some individuals from freshwater to marine habitat over multiple years [51,88]. Our results generally corroborate this but also provide evidence for increased variation in occupied habitats within anadromous types than previously documented in Broad Whitefish. We found major differences in the duration of freshwater and estuarine use prior to marine migration, which ranged from as little as a month or so to several years. In other fish species, juvenile life history diversity appears to be driven by differences in habitat quality and availability and the response of individuals to maximize fitness [105,106]. Chinook Salmon, for example, exhibit a continuum of juvenile life history pathways, which are defined by the timing and transition among life stages and freshwater and marine habitats used for rearing [106]. These life history pathways are influenced by local habitat and environmental conditions, such as water temperature, which influences factors such as incubation duration, growth rate, and downstream migration [106]. In many instances, Chinook Salmon that are exposed in warmer post-emergence water temperature display higher growth rates, which generally results in earlier downstream migration [107]. Similar processes along with genetically-controlled traits, such as adult fish spawn timing and adaptation to spawning environments, likely influence the diversity of Broad Whitefish life histories documented here.

Semi-anadromous life history types occurred within our samples of Colville River Broad Whitefish but, to our surprise, the vast majority of these individuals did not exhibit a freshwater natal signature, suggesting they had been advected into estuarine and or marine environments soon after hatching. Subsequently, these individuals frequently moved between estuarine and marine habitats throughout their lives, with minimal time spent in fresh water. This pattern suggests that adult adaptation to spawning environments near coastal estuarine and marine habitats may influence life history trajectory [108], as seen in other fish species such as Atlantic Salmon (*Salmo salar*) and Brown Trout (*S. trutta*) [109]. Semi-anadromous individuals were the most common life history type documented, which suggests that nearshore and marine habitats are significantly important for juvenile Broad Whitefish born in the Colville River.

Nonanadromous Broad Whitefish have also been documented in the Yukon and Mackenzie rivers [51,38,102]. In the Yukon River drainage, nonanadromous individuals are less common than anadromous individuals [51], with the exception of sites far inland (i.e., 2000 km from the coast) and in upstream tributaries like the Porcupine River. Similar patterns have been observed in the Mackenzie River drainage, where nonanadromous individuals are uncommon and tend to be associated with lentic habitat [38,103] or smaller tributaries (e.g., Arctic Red River) [104]. Our results show a similar pattern in the Colville River watershed but provide evidence for additional diversity within nonanadromous life history types. The observed habitat use of freshwater estuarine nonanadromous life history types suggests that certain individuals do not remain in one location and rather move between a variety of freshwater and estuarine habitats. Collectively, these findings suggest that migrating to sea is not energetically profitable for all Broad Whitefish, such as those requiring particularly long journeys, and that enough resources to reach sexual maturity can be acquired solely in fresh water.

### Age-0 movements

The majority of Broad Whitefish appear to be transported from natal habitat to rearing habitat within the first year of life. Generally, otoliths indicated that individuals with life history types 1–2 and 5–6 left natal habitat and utilized freshwater rearing habitat with isotopically different values. Our results suggest that most larvae do not use natal habitat for rearing and instead are likely transported downstream to new habitat. In contrast to previous research, we found that numerous individuals (n = 13) entered marine habitat at age-0 after spending little time in freshwater. To our knowledge, the rapid transition over days instead of weeks has not been documented but might be due to the proximity of spawning habitat to marine environments and the intensity of spring breakup streamflow on certain years. Across numerous freshwater fish species, there is evidence that spawning areas tend to be associated with shallow habitats [115] and provide refugia for eggs where instream conditions facilitate embryonic development and predation is minimal [116]. For example, salmon and trout commonly use small streams or braided channel networks to spawn, taking advantage of habitat that often contains high hyporheic flow, consistent water temperatures, and gravel substrate [117,118,119]. Within heterogenous riverine environments, it is common for spawning habitats to differ from juvenile rearing habitats, which may also be distinct from adult habitats [44,45]. These patterns of habitat use has been documented for Broad Whitefish in the Yukon and Kuskokwim river drainages, where adults feed in lakes during the summer, spawn in upper reaches of the drainage during the fall, and larvae and juveniles rear in downstream delta or estuary habitat [49, 50, 52].

A subset of individuals exhibited minimal isotopic change between natal and juvenile rearing regions. The late transitional anadromous life history type (Type 3), for example, displayed minimal isotopic change, which could be caused by individuals being advected downstream to new rearing habitat that were isotopically similar prior to marine entry. The semi-anadromous type (Type 4) also had minimal isotopic change and, in this case, it is possible that they are not advected far or end up rearing in nearby freshwater habitat with isotopic values similar to natal areas (e.g., lakes, riverine habitat).

Identifying freshwater natal and juvenile rearing specific habitat is difficult due to the lack of empirical ^87^Sr/^86^Sr data, but several broad patterns emerged from comparing our results to coarse estimates of surface water ^87^Sr/^86^Sr (1 km^2^ grids) for surface waters across Alaska [68,69]. Broad Whitefish are known to spawn in wide channels with moderate braiding and gravel substrate [49,50,88]. Recent research in the Colville River estimates mainstem habitat to have the greatest intrinsic spawning potential [78], which is where we would expect natal habitat to be located, as opposed to smaller tributaries with similar ^87^Sr/^86^Sr. Anadromous individuals that migrated to sea at age-0 (Type 1) had natal region ^87^Sr/^86^Sr that were higher than juvenile freshwater rearing region ^87^Sr/^86^Sr, suggesting that individuals were born in the middle watershed where estimated ^87^Sr/^86^Sr is near 0.711 and then gradually decreases downstream.

Late migrating anadromous individuals (Type 2–3) had juvenile region ^87^Sr/^86^Sr that were higher than natal regions. This pattern could be explained by individuals hatching from eggs spawned higher in the watershed, where the estimated ^87^Sr/^86^Sr range was near 0.710, drifting downstream to rear in habitats in the middle and lower watershed where estimated ^87^Sr/^86^Sr was generally higher (ca. 0.711). Semi-anadromous individuals (Type 4) had natal and juvenile region ^87^Sr/^86^Sr near the GMV (0.70918), which suggests they moved between areas in the lower river, delta, and estuary. Nonanadromous individuals (Type 5–6) had higher natal region and lower juvenile rearing values, suggesting they may have hatched in the middle watershed drifted downstream to rear in the lower river or estuary, or in the case of the freshwater resident life history type, remained in the spawning area.

### Broad Whitefish conservation

Our research provides new insights into the complex patterns of lifetime habitat use by Broad Whitefish. A portfolio of life history types suggests that Colville River Broad Whitefish largely remain free from anthropogenic impacts that fragment or homogenize habitats. Within life history types, individuals appear to use different habitats (freshwater, estuarine, and marine) for varying durations (months to years) across each life stage, which suggests further complexity. For example, an individual may hatch and spend less than one month or as much as five years in freshwater before heading to sea. Once at sea, some individuals continue annual patterns of marine migrations through life, while others migrate as mature adults to freshwater habitats they have not previously occupied. There is certainly greater life history variation than the small number of otolith ^87^Sr/^86^Sr profiles examined here would suggest. Similar diverse patterns of habitat use have been documented for other fish species. Recent otolith microchemistry work with juvenile Chinook salmon, for example, demonstrated that individuals used a complex array of habitat types to achieve maximum growth prior to ocean migration [120]. The diversity of life histories found here suggests that Broad Whitefish populations experience frequent disturbance or high environmental variability [100,108,113].

Our research revealed the complexities of Broad Whitefish life histories and habitat use, but further research is needed for the effective conservation of this important subsistence resource. Understanding the variation in ^87^Sr/^86^Sr across the watershed is important for identifying spawning habitats, which are essential to sustain Broad Whitefish populations and may be disproportionally small [78] compared to other key habitats (e.g., feeding habitat). Once identified, it will also be possible to understand the contribution of each spawning area to the Nuiqsut subsistence fishery and potentially to other fisheries [62]. Broad Whitefish are a highly migratory and long-lived fish, making it conceivable that fish caught outside the watershed (e.g., in the Utqiagvik, AK, fishery), may have originated from the Colville River. Lastly, another challenge is understanding where fish go when they leave the Colville River watershed. Future research that tracks the marine migration patterns and movement between watersheds across the Arctic will likely provide key insights into a diversity of juvenile and adult foraging patterns and migration routes.

Despite its historical and current importance as a subsistence species for Alaska’s Indigenous Beaufort Sea communities, little is known about the nature and distribution of their essential habitats. Currently, infrastructure from oil and gas development is minimal within the Colville watershed. However, as development continues to expand across the region, potentially crossing and altering habitats in the Colville River and other watersheds utilized by Broad Whitefish, it is important for land managers and conservation planners to understand the risks.

Oil and gas development in the central Beaufort Sea region has caused cumulative impacts to permafrost, such as flooding and pooling of water, and loss of vegetation due to heavy road dust [121,122], which can cause flow modifications that affect habitat quality and connectivity. Arctic oil and gas infrastructure (e.g., roads, pipelines) fragments and disrupts aquatic ecosystems along linear paths [126], which can further introduce stressors to juvenile and adult fishes [25,123], such as increased sedimentation [124–127], modifications of streamflow [128], obstructions to passage [129–131], and reduced instream habitat quality [132], as well as pollution [133]. Climate change is also altering Arctic hydrologic regimes; variability in runoff is increasing [14], and discharge from large Arctic rivers is increasing both annually [15,16] and during winter [17], and snow-dominated runoff regimes are shifting toward rainfall-dominated regimes [134]. Maintaining habitats to support complex life histories is critical for the long-term conservation of Broad Whitefish and the loss of life history diversity will make the population more susceptible to fluctuations in abundance and will increase the risk of extinction [135–137].

## Acknowledgments

We would like to thank the Alaska Cooperative Fish and Wildlife Research Unit (AKCFWRU) staff, Native Village of Nuiqsut (NVN) Tribal Council staff, U.S. Fish and Wildlife Service Fairbanks Fish and Aquatic Conservation Office staff, Bureau of Land Management (BLM) Arctic Office staff, and the Alaska Established Program to Stimulate Competitive Research for assistance with fieldwork and logistics. We also thank Jonah Nukapigak (NVN) for crucial field logistical support and for sharing his traditional knowledge, and Stuart Lehman, Mike Lunde (AKCFWRU), and Kristin Rine (AKCFWRU) for critical assistance in the field with Broad Whitefish capture. We also thank Lonnie Bryant (BLM) for assistance with summer activity permits and Richard Kemnitz (formerly BLM) for logistical help in Umiat, AK. We thank Josh Gage with Gage Cartographic for assistance with R code to help organize and analyze strontium otolith data. We thank Andrew C. Seitz (CFOS, UAF) and Jeffrey A. Falke (USGS AKCFWRU) for reviewing early manuscript drafts and providing constructive comments. Research was conducted under the Alaska Department of Fish and Game Fish Resource Permit #SF2017-201 and under University of Alaska Fairbanks Institutional Animal Care and Use Committee Protocol #901048. The findings and conclusions in this article are those of the authors and do not necessarily represent the views of the U.S. Fish and Wildlife Service. Any use of trade, firm, or product names is for descriptive purposes only and does not imply endorsement by the U.S. Government.

## Author contributions

JCL^1,2^, DJR^3^, MSW^4^, RJB^5^, and MSW^7^ conceived of or designed the study; JCL^1,2^, RJB^5^, and KJS^6^ developed methods; JCL^1,2^ performed field research, JCL^1,2^, RJB^5^, and KJS^6^ conducted lab research; JCL^1,2^, DJR^3^, RJB^5^, and KJS^6^ analyzed data with support (review and critical feedback) from MSW^4^ and MSW^7^; manuscript preparation was led by JCL^1,2^, with support (critical feedback, revisions, and additions to text) from all co-authors.

## Supporting information

**S1 Table. Summary of Broad Whitefish sampled.** Table displaying population sex, age, and size structure, tissues sampled, and sample size for Broad Whitefish (*Coregonus nasus*) caught at three locations within the Colville River watershed, Alaska, USA.

**S2 Table. Laser Ablation System (LA) multi collector inductively coupled plasma mass spectrometer (MC-ICP-MS) instrument parameters.** Table displaying the LA MC-ICP-MS instrument parameters used at the Alaska Stable Isotope Facility, University of Alaska Fairbanks, Fairbanks, AK, USA.

**S3 Table. Summary of Broad Whitefish (*Coregonus nasus*) sampled in the Colville River, AK, USA.** Table displaying otoliths analyzed and attributes.

**S1 Figure. Sr otolith plots.** Plots showing ^88^Sr concentration and ^87^Sr/^86^Sr for all Broad Whitefish (*Coregonus nasus*) caught within the Colville River watershed, Alaska, USA. We cropped strontium data for otoliths analyzed (n = 61) at the otolith core and edge.

